# Climate and genetic diversity change in mammals during the Late Quaternary

**DOI:** 10.1101/2021.03.05.433883

**Authors:** Spyros Theodoridis, Alexander Flórez-Rodríguez, Ditte M. Truelsen, Konstantinos Giampoudakis, Raquel A. Garcia, Joy Singarayer, Paul Valdes, Carsten Rahbek, Katharine A. Marske, David Nogués-Bravo

## Abstract

Conservation decisions and future scenarios are in need of past baselines on climate change impacts in biodiversity. Although we know that climate change has contributed to diversity shifts in some mammals^1,2,3,4,5,6,7^, previous research often assumed that climate change is invariable across species’ ranges. We are therefore still ignorant of the true rates of climate change experienced by species assemblages over the last millennia, their impacts on intraspecific diversity, and how they compare to future climate change projections. Here, we use more than 9,000 Late Quaternary records, including fossils and ancient and modern DNA sequences, millennial-scale paleoclimatic reconstructions over the last 50,000 years and future climate change projections to document rates of climate change velocity and dynamics in genetic diversity experienced by an assemblage of 16 extinct and extant Holarctic mammal species. Extinct megafauna experienced velocities more than 15 times faster than the extant species, up to 15.2 km per decade. Notably, extant large-bodied grazers lost almost a 65% of their pool of genetic diversity since the Late Pleistocene, which indicates reduced ability to adapt to on-going global change. Additionally, mammal species experienced overall climate change velocities slower than that projected for the end of the 21^st^ century but punctuated by comparable fast climate change episodes. Our results provide baselines on the impacts of ongoing and future climate change on the diversity of mammal species.

## Main

As we enter into a period of accelerating climate change and elevated extinction rates^8^, identifying the magnitude of past episodes of climate change experienced by species and its impacts on their biological diversity is vital to anticipating and mitigating both species extinction^9, 10^ and ecosystem degradation^10, 11^. Over the last 50,000 years climate change have triggered substantial population declines, extirpations and extinctions^1, 3^. These past climatic footprints in biodiversity can provide baselines on the extent to which animal and plant species were subject to environmental changes^12^. The rapid accumulation of paleoclimatic data, fossil and ancient DNA data has opened the door to assessing species demographic fluctuations through the Late Quaternary climate changes^5–7^. Yet, little is known about rates of past climate change actually experienced by extinct and extant mammal species across large regions of the planet, and the extent to which these rates may have driven the erosion of intraspecific genetic diversity, a fundamental component for species adaptation and persistence under changing environments^13, 14^. Previous studies have examined whether fluctuations in species genetic diversity and demography are concomitant with past climate events, but generally assumed that past climate trends extrapolated from a single location, such as Arctic or Antarctic ice cores, reflected the local climatic conditions experienced by species across their geographic distributions^3, 6, 15^. This limitation has precluded a deeper understanding of biodiversity shifts and loss under climate change.

Here, we investigate the velocity of climate change and changes in intra-specific genetic diversity over the last 50,000 years across 16 terrestrial mammals (plus two marine mammals without available climate information) with different life history and ecological traits. We compiled 9,100 georeferenced paleorecords from the Late Pleistocene (between 50,000 and 11,700 y BP) and the Holocene epochs (11,700 y BP to present) (Supplementary Table 1), including 5,780 radiocarbon-dated fossils, 1,142 aDNA and 2,178 modern mitochondrial DNA sequences (Cytochrome b and Control region, Fig. 1a). We spatially matched the records fulfilling quality control criteria (Methods) to paleoclimatic reconstructions (1×1 degree resolution) extending back 50,000 years^16^ (at a temporal resolution of 1-2 millennia) and estimated the climate change velocity^17^ (km/y) experienced by each species across its range for 36 time periods. We used the ancient and modern DNA sequences to estimate genetic diversity (that is, haplotype and nucleotide diversity) for each species between the Late Pleistocene and the Holocene (before and after 11,700 years ago), and further estimated changes in genetic diversity before and after the fastest and slowest climate change velocity event experienced by each species. Finally, we tested whether variation in the estimated changes in genetic diversity within the assemblage of mammal species can be explained by climate change velocity or whether changes were species specific.

**Fig 1.**
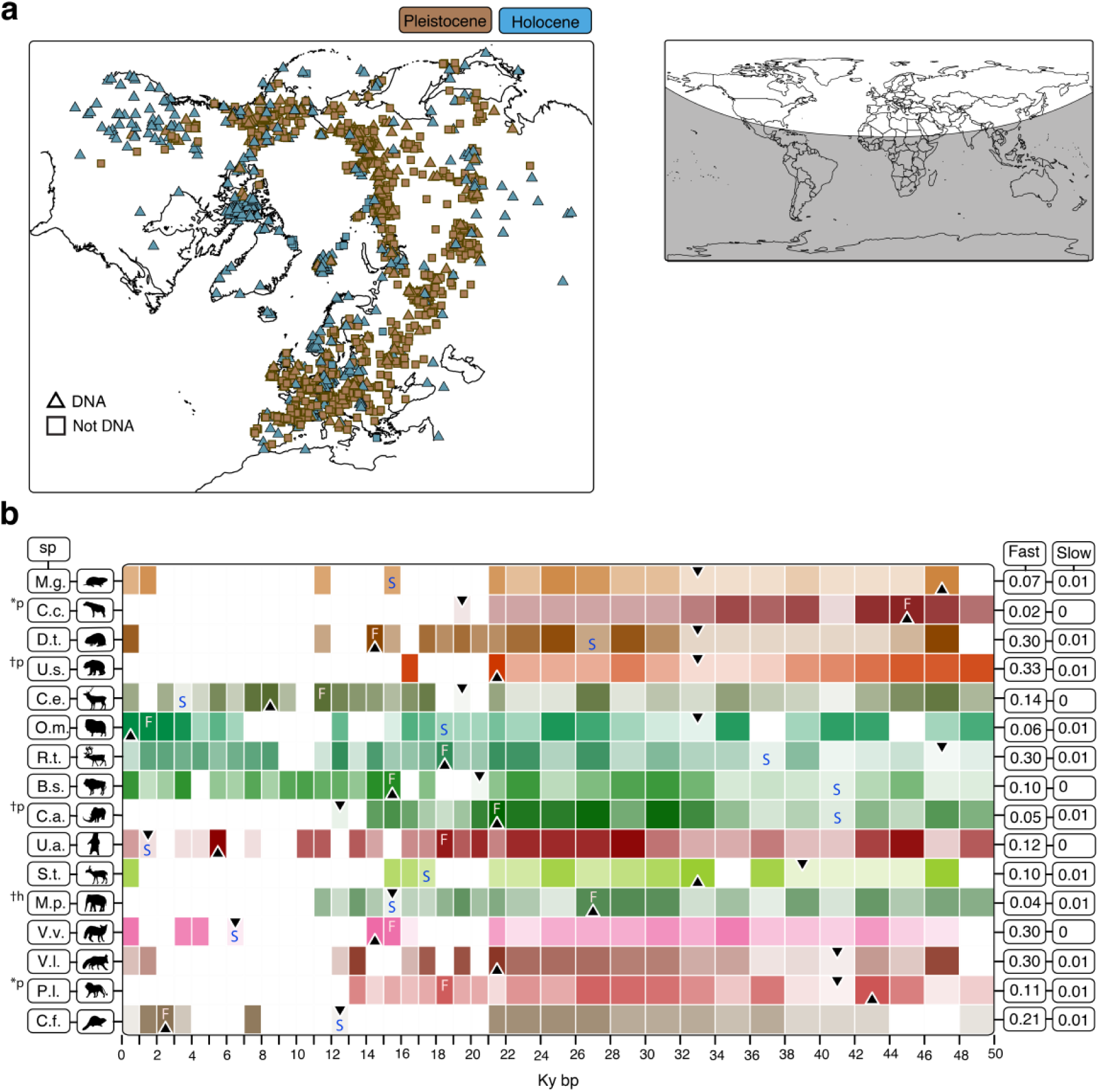
Distribution of paleorecords and climate change velocity through time. (**a**) DNA sequences and radiocarbon-dated fossils for 18 Holarctic mammals. White area in the world map in the top-right depicts the study area. (**b**) Climate change velocity experienced by 16 Holarctic terrestrial mammals during the last 50,000 years. Colored squares represent species’ median climate change velocity (km per year) at each millennial or bimillennial period, with darker colors representing faster velocity. Blank squares represent periods with no records. Black up- and down-pointing triangles indicate the millennial periods with the fastest and slowest climate change velocity for each species, respectively. The extreme values are indicated in the boxes at the right of the figure. Letters F (white) and S (blue) indicate the fastest and slowest millennial periods for each species, respectively, used in the associations with genetic diversity change. Species are ordered from the fastest (bottom) to the slowest (top) median climate change experienced during the last 50,000 years. C.f.= *Castor fiber* (Eurasian beaver), P.l.= *Panthera leo* (lion), V.l.= *Vulpes lagopus* (arctic fox), V.v.= *Vulpes vulpes* (red fox), M.p.= *Mammuthus primigenius* (woolly mammoth), S.t.= *Saiga tatarica* (saiga antelope), U.a.= *Ursus arctos* (brown bear), C.a.= *Coelodonta antiquitatis* (woolly rhino), B.s.= *Bison sp.* (bison), R.t.= *Rangifer tarandus* (reindeer), O.m.= *Ovibos moschatus* (musk ox), C.e.= *Cervus elaphus* (red deer), U.s.= *Ursus spelaeus* (cave bear), D.t.= *Dicrostonyx torquatus* (arctic lemming), C.c.= *Crocuta crocuta* (cave hyena), M.g.= *Microtus gregalis* (narrow-headed vole). †p = extinct in the Pleistocene, †h = extinct in the Holocene. *p = Holarctic extirpation in the Pleistocene.

Climate change velocity experienced by Holarctic mammals varied considerably among taxa as well as within taxa across time over the last 50,000 years (Fig. 1b, Supplementary Table 2). Climate change velocity ranged from 0.0001 to 1.52 km/y for all terrestrial fossil and DNA localities (*n* = 8656), with 50% of those localities experiencing velocities slower than 0.023 km/y. Remarkably, the extinct megafauna, including woolly mammoths (*Mammuthus primigenius*), woolly rhinos (*Coelodonta antiquitatis*), cave bears (*Ursus spelaeus*) and cave hyenas (*Crocuta crocuta*), experienced climate change velocities at least 15 times faster (0.344 to 1.52 km/y) than any other extant Holarctic species (Fig. 2, Supplementary Table 2), suggesting a major role of past climate change on species extinction.

**Fig 2.**
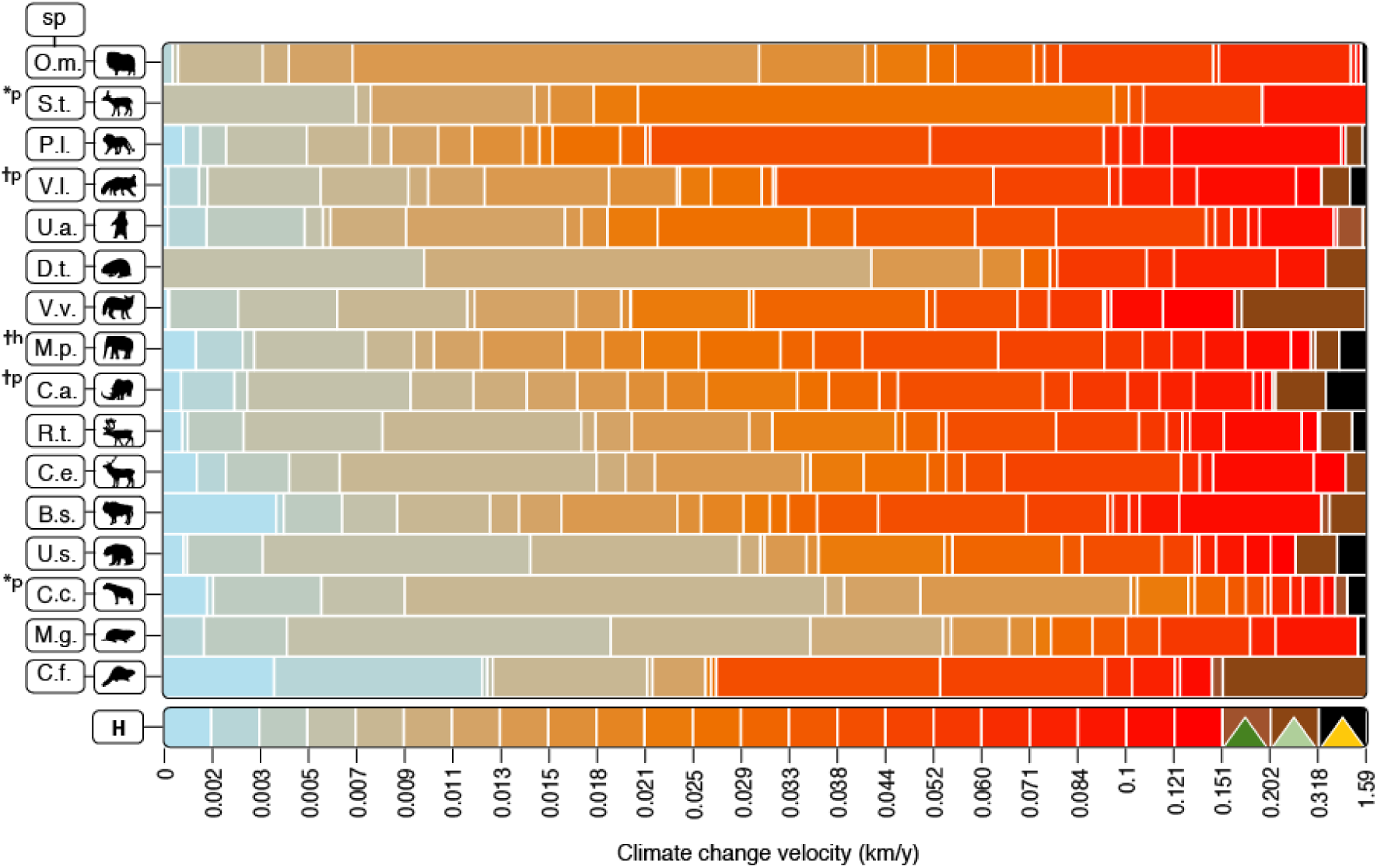
Climate change velocity experienced by individual species relative to the whole Holarctic region. The bottom row indicates the range of climate change velocities detected across the Holarctic during the last 50,000 years, shown as equally sized rectangles ranging from blue (slow) to red (fast). Horizontal bars for each species indicate the proportion of grid cells (that is, cells with paleorecords and modern samples) experiencing different velocities relative to the total number of grid cells through time for the respective species. For example, the relative proportion of grid cells within the lowest two magnitudes of climate change velocity (light blue) for *Castor fiber* (bottom species row) is higher than any other species. Green, mint and yellow triangles in the bottom right represent predicted climate change velocity for the year 2100 for temperate, polar and cold regions^17^ respectively. The most recent records for each species were used to estimate exposure to future climate change velocity.

Changes in genetic diversity from the colder and climatically less stable Late Pleistocene to a warmer and more stable Holocene varied considerably among species (Fig. 3, Supplementary Fig. 1). Two-thirds of all species lost nucleotide diversity, ranging from comparatively small reductions (e.g., 8% for reindeer) to global extinctions before the Holocene. Notably, ten out of the twelve species that lost genetic diversity were megafauna (>44 kg), mainly grazers. Among the six species with higher nucleotide diversity in the Holocene than in the Late Pleistocene, four weigh less than 30 kg, including the red (*Vulpes Vulpes*) and arctic (*Vulpes lagopus*) foxes, the narrow-headed vole (*Microtus gregalis*) and the Eurasian beaver (*Castor fiber*) (Fig. 3, Supplementary Fig. 1). Contrasting patterns were detected among the marine mammals, with the bowhead whale (*Balaena mysticetus*) losing and the grey whale (*Eschrichtius robustus*) gaining genetic diversity. Changes in genetic diversity between the Late Pleistocene and Holocene do not seem to be related to different sample sizes between these two epochs (Supplementary Fig. 2 and 3).

**Fig 3.**
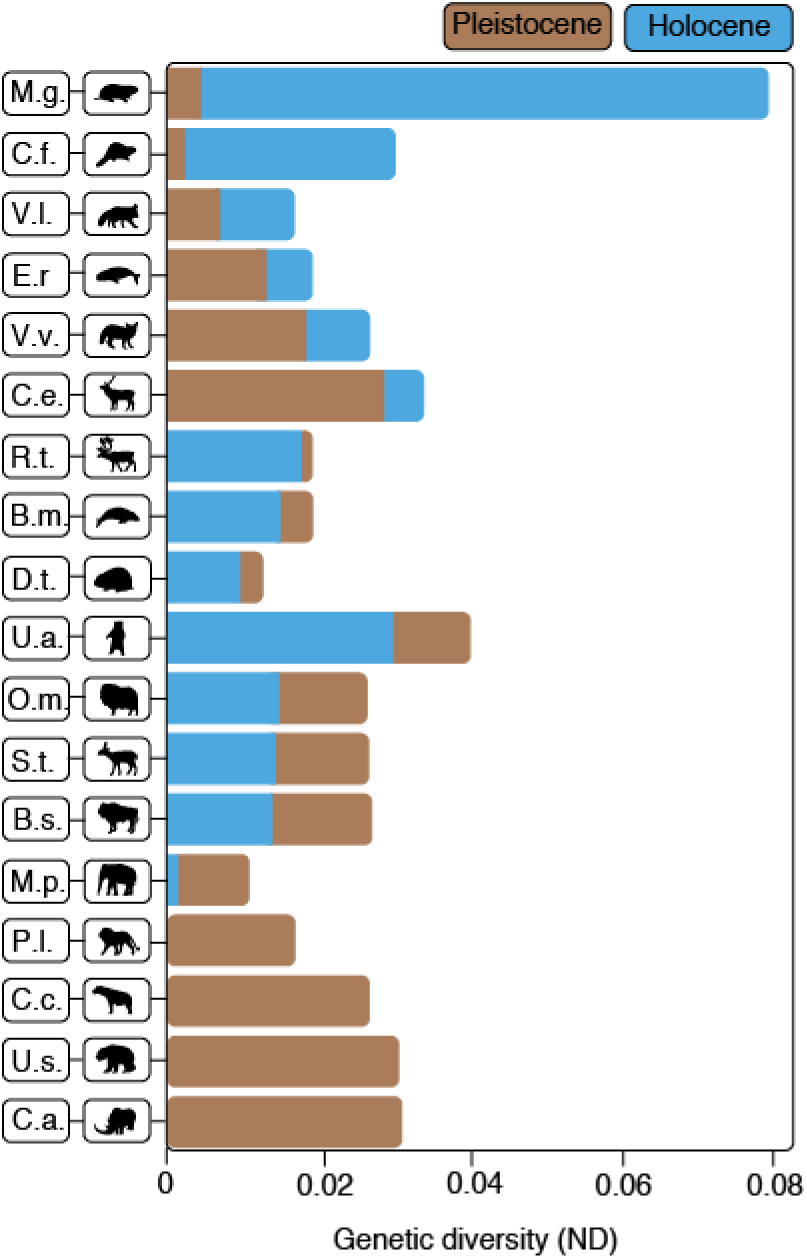
Pleistocene and Holocene genetic diversity per species. Nucleotide diversity (ND) in Pleistocene (brown) and Holocene (blue) for each of the 18 Holarctic mammals are shown as overlapping bars. Species are sorted from bottom to top by percentage of ND loss between the Pleistocene and Holocene. Species’ name abbreviations as in Fig. 1, with the addition of B.m. = *Balaena mysticetus* (bowhead whale), E.r.= *Eschrichtius robustus* (grey whale).

During the Late Quaternary the Holarctic region witnessed a decline in mammal species richness^18^ and floral diversity^19^, including the extirpation and replacement of major genetic lineages^3, 20^. Large-bodied grazers with long generation times now have approximately half of the nucleotide diversity (56%, sd = 41%; Fig. 3) they had in the Late Pleistocene. The transition from a more genetically diverse Late Pleistocene to a genetically eroded Holocene, together with the expected positive correlation between regional species richness and genetic diversity^20, 21^, suggest linked responses across ecosystems, species and genes. For instance, across the Holarctic, the replacement of dry steppe-tundra by moist tundra ecosystems may have reduced the carrying capacity of the environment for herbivorous megafaunal species^19^. These changes led to local extinctions, range contractions and range shifts, all of which would have resulted in decreased levels of intraspecific genetic diversity^22^. However, other species thrived, and many of those mammals that harbor more genetic diversity today than they did in the past (Fig. 3) share traits like short generation time or high fecundity, highlighting how life history traits affect a species’ ability to persist under global environmental change^23^.

Whether climate change, humans or a combination of both caused the extinctions of megafauna species remains a heated debate. However, several lines of evidence suggest that climatic changes during the Late Quaternary prompted the turnover of megafaunal assemblages^3^, increased the likelihood of global extinctions^24^, and shaped the distributions and regional richness of endemic species^25^. These dynamics reflect species’ responses, often idiosyncratic, to past climate change, such as local extirpations, contraction into refugia or range shifts tracking suitable environments^5, 26^. Millennial-scale climate change velocity in the Late Quaternary does not explain changes in haplotype diversity (Fig. 4a; *R*^2^ = 0.01, P = 0.642) or nucleotide diversity (Fig. 4b; *R*^2^ = 0.001, P= 0.867) across species, suggesting that climate velocity likely did not drive an assemblage-wide level effect in genetic diversity of mammal assemblages. Dissimilarities in species’ climatic tolerance, dispersal ability and other traits likely led to species-specific responses, yielding both winners and losers under past climate change.

**Fig 4.**
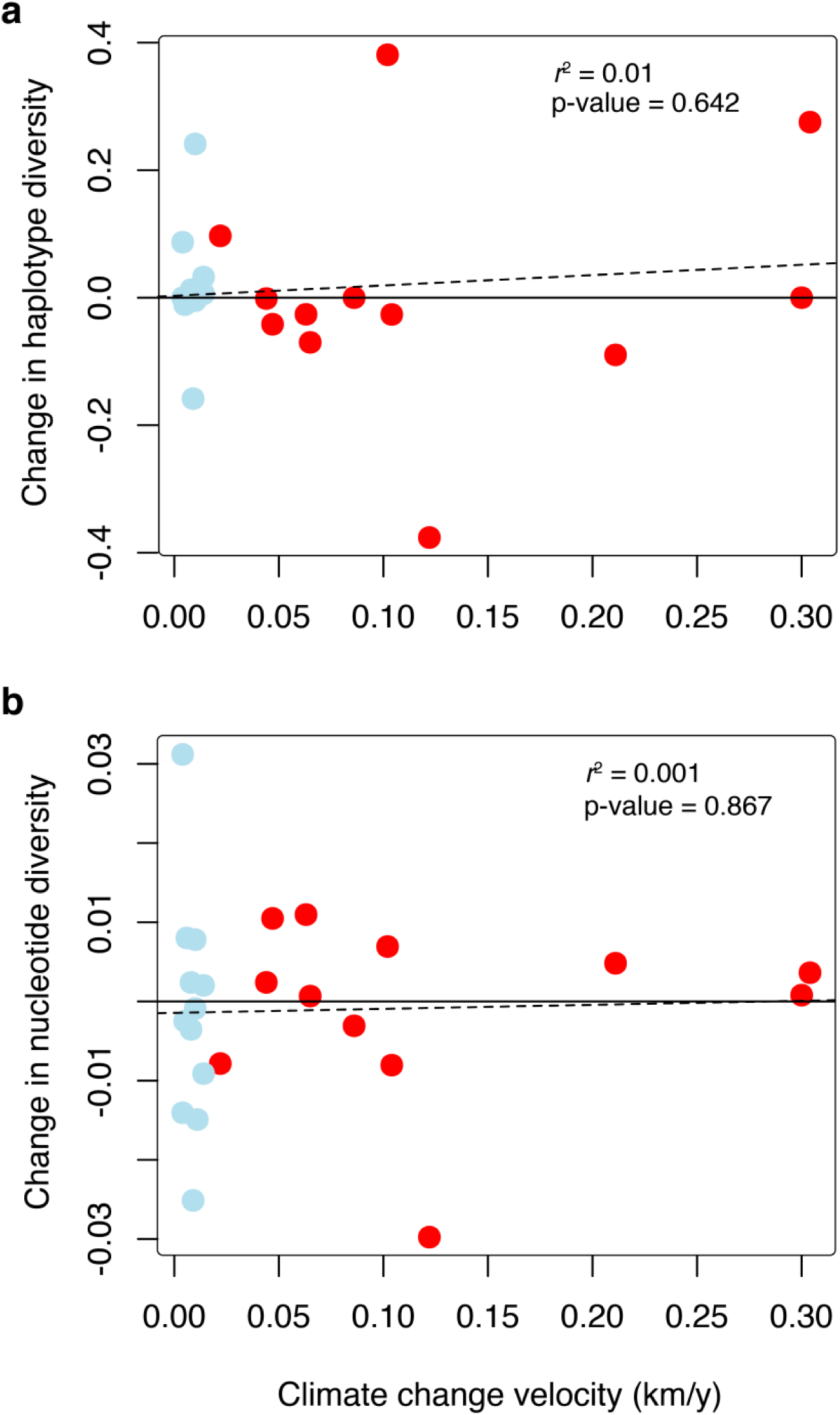
Correlations between climate change velocity and change in genetic diversity. Correlations between the fastest (red) and slowest (blue) median climate change velocity experienced by each species (Fig. 1b) and changes in Haplotype diversity (**a**), and Nucleotide diversity (**b**) for each species. Changes in genetic diversity were estimated using bins of 5,000 years before and after the selected time period of climate change velocity (see Methods). Dotted line shows the linear model for the variables.

Between now and the end of the 21^st^ century, species are expected to face climate change velocities that significantly exceed even the fastest velocities experienced during the last 50,000 years^27^ (Fig. 2 and Supplementary Fig 4). Considering each millennial period separately, median climate change velocity^18^ across the Holarctic reached the fastest rates, 0.18 km/y, 11,000 years ago (Supplementary Fig 4), which is noticeably slower than the projected velocity for temperate (0.2 km/y), polar (0.25 km/y) and cold (0.45 km/y) regions by 2100 (Fig. 2). Given that our species experienced velocities during specific periods of time similar or slower than those projected for the year 2100 (Fig. 2), we can expect increasing challenges for species, particularly for those species with large body sizes and long generation times. Our estimates of past climate change velocity are conservative in relation to future projections, as they are calculated between millennial time periods, and they do not capture past events of abrupt and fast climate change at the decadal time scale^28^.

Past “natural experiments” of biodiversity shift under climate change hold great potential to provide baselines and insights on current and future biological responses to climate change. With species likely to experience faster climate change than in the past, the potential future loss of populations due to fast climatic shifts over space may trigger decreases in the overall genetic pool of large body-size mammals, as reported here for some species, heightening concerns about their future conservation status^29^. Moreover, anthropogenic factors such as land-use change, invasive species or over-harvesting are also causing local population extinctions and species range contractions^30^, further contributing to the decline of biological diversity.

## Methods

### Database assembly

#### Species selection and DNA sequence retrieval

To evaluate the impacts of climate change on genetic diversity throughout the Late Quaternary, we compiled the most comprehensive (to date) comparative database of Holarctic mammals with publicly available ancient DNA (aDNA), modern DNA and radiocarbon-dated fossils. We restricted our study area to the Holarctic because it encompasses the densest and highest quality fossil and aDNA record across biogeographic regions. Moreover, the Holarctic region experienced intense climate change, including phases of warming and cooling. First, we conducted a literature search using the key words: [“aDNA” OR “ancient DNA” OR “historical DNA”] in ISI Web of Science. The search was conducted in December 2014 and yielded 6,618 studies. From these results, we narrowed our search to those with mitochondrial sequences in both Late Pleistocene (LP, from 50,000 to 11,700 years BP) and Holocene (from 11,699 years BP to present). We omitted species for which domestication may have had a significant impact on intraspecific genetic variation, including *Equus caballus*, *Canis lupus* and *Sus scrofa*. For the remaining 18 species (Supplementary Table 1), we downloaded all mitochondrial sequences available in GenBank (accessed in July 2015), including aDNA and modern DNA sequences. As *Panthera leo* and *Crocuta crocuta* are extinct in the Holarctic region, only aDNA sequences were included for these species. To generate comparable estimates of genetic diversity, we retained sequences from the loci with the greatest number of aDNA sequences available for each species, and used the same genetic marker for the Late Pleistocene and Holocene populations. We chose *cytochrome b* and *control region* as focal loci, as these markers have been widely used in previous aDNA studies. Sequences from zoological gardens and/or animal parks were also excluded. For all the remaining aDNA sequences, dating information was extracted from the original studies when available. In total, our database comprised 4,794 DNA sequences (756 from the Late Pleistocene and 4,038 from the Holocene) for 18 species, including three extinct species. It represented six orders and 12 families of mammals with Holarctic distributions during at least part of the Late Pleistocene (Supplementary Table 1).

#### Retrieval of fossil occurrences and calibration of ^14^C dates

We compiled a fossil database of mammal species that were present in the Holarctic during at least some part of the terminal Late Pleistocene and Holocene transition, and that also met the DNA sequence requirements as specified above (18 species in total). We used the publicly available Stage 3 Project database^31^ and selected publications of aDNA studies and of past species distributions as source information for the species occurrences. A total of 6,971 radiocarbon-dated samples (5,780 fossil and 1,191 aDNA) that span from 50 ky ago to the present were included in our study (Fig. 1a). The majority of these samples were directly dated using a variety of techniques that rely on some physical or chemical property of the sample (i.e. radiocarbon ^14^C), thermoluminescence (TL), uranium-series (U-Th), or electron-spin resonance (ESR) dating). Of these, the vast majority were ^14^C dated - both by beta counting ^14^C and Accelerator Mass Spectrometry (AMS). We also used some indirectly dated samples found in the same site and stratigraphic level as directly dated specimens that were determined genetically or morphologically to belong to different taxa. These were assigned the same age determination and error as the directly dated specimens.

We used the indirectly dated records to compare genetic patterns between the Late Pleistocene and Holocene (Fig. 3) as the time span of these glacial epochs is much wider than the dating uncertainty of the specimens used. However, for the climate change velocity analysis (Fig. 1b and Fig. 2) we used smaller time bins (1,000 and 2,000 years), which increased the likelihood that some occurrences might have a higher dating error than the time bin span (Supplementary Fig 5). Thus, only the fossils with ^14^C-direct and indirect (as defined above) radiocarbon determinations were used for climate change velocity analysis. To further increase the quality of our database we also excluded: 1) samples with duplicate laboratory codes, 2) samples with more than one age determination, 3) samples described as contaminated by the source information and 4) samples lacking laboratory codes or coordinates, with uncalibrated ^14^ C dates or ^14^ C dating error (sigma). Radiocarbon dates for both aDNA and fossils were calibrated to calendar years using the OxCal v4.2 program and the IntCal13 radiocarbon calibration curve^32–34^.

#### Georeferencing

To integrate our fossil, aDNA and modern DNA occurrences with spatially explicit paleo-climatic simulations, we obtained geographical coordinates for all terrestrial fossil and DNA samples using information listed in GenBank, Stage 3 Project fossil database and the primary literature. However, when the original source lacked coordinates but provided locality descriptions, we assigned the coordinates using the API tool provided by GeoNames.org^35^. This tool assigns latitude and longitude to locality descriptions and has recently been used to map thousands of genetic sequences from the GenBank and BOLD databases^36^. When the locality was not available through GeoNames.org, we conducted internet-based literature searches together with the Google Maps spatial search engine. Both GeoNames.org and Google Maps rely heavily on the quality of the locality description, and for many specimens, locality descriptions were relatively broad (e.g., Eastern Canada). To identify the level of geographic precision among the coordinates of all the occurrences in the database, we ranked them as follows: 1) coordinates available from the original source (*n* = 6,793, 71%); 2) locality description either with coordinates from other studies or location available in Geonames.org or Google Maps (*n* = 1,083, 11%); 3) broad locality description such as mountain range, national park, country region (*n* = 1,224, 13%); and 4) only country information (*n* = 472, 5%). Records with rank 4 were excluded from the analyses.

### Climate change velocity

#### Paleoclimatic data

Paleoclimatic simulations were used to estimate the velocity of climate change across the last 50,000 years. The paleoclimatic reconstructions are based on the Atmospheric-Ocean coupled General Circulation Models (AOGCMs) that simulate the coupled components of the Earth’s climate system, taking into account the interactions between the atmosphere, oceans, land surface and ice sheets^37^. These simulations are used to predict past, present and future climate changes^38^. The paleoclimate simulations used in this study were constructed with the HadCM3 (Hadley Centre Coupled Model version 3) AOGCM^39^. The paleoclimate simulations have been reassembled to a resolution of 1×1 degrees^16^. Climatic simulations are bimillennial between 50 kyr BP and 22 kyr BP and millennial from 22 kyr BP to present, yielding 36 paleoclimatic reconstructions in total.

#### Climate change velocity

Species’ ability to track suitable climates is constrained by the magnitude of climate change and the distance required to track constant climatic conditions. Therefore, we used climate change velocity as the measure of climate change to compare to changes in genetic diversity. Climate change velocity is a measure of the pace at which climatic conditions move across the local topography^17, 25, 27, 40, 41^. Climate change velocity is an imperfect measure of exposure to climate change-related population declines or extinctions, as it underestimates exposure in mountain regions^42^. However, the geographical extent (and resolution) of our study is covered mainly by large plains across central and north Eurasia and is lacking large mountainous regions as the Himalayas or the Andes. It thus offers a study system that minimizes the key limitation of the climate velocity estimate. As paleoclimatic reconstructions for precipitation have a higher level of uncertainty than those for temperature^25^, we used mean annual temperature to estimate climate change velocity. Following previous studies^17, 25, 27^, we calculated climate change velocity by dividing the temporal temperature gradient (difference in °C between the focal and the preceding time bin) to the spatial climate gradient (°C / km), where the spatial climate gradient is a measure of the variability in temperature between adjacent grid cells (3×3 grid cell neighbourhood) in the preceding time bin. The temporal temperature gradient was divided by the length of the time bin (either 1,000 or 2,000 years) to obtain temperature change per year. To avoid dividing the temperature change per year by a spatial variability of zero or values close to zero, values for the spatial variability below 0.00005 °C km^1^ were set to be equal to 0.00005 °C km^-1^ ^17, 25^. Climate change velocity is measured in °C yr^-1^/°C km^-1^ = km yr^-1^. Values for projected climate change velocity between 2100 and present time were taken from a previous study^17^. We then spatially and temporarily matched the filtered paleorecords for each of the 16 terrestrial mammals to the climate change velocity estimates for each time bin. We finally calculated the median climate change velocity per species and time bin (Fig. 1b and Fig. 2). All calculations were performed in R version 3.01^43^ using the “raster” package^44^. Climate change velocity for the Holarctic region is shown in Supplementary Fig 4.

### aDNA sequence preparation

#### Sequence alignment

We performed DNA sequence alignments for all the species in the database using MUSCLE^45^ with default settings as implemented in the program Geneious R8^46^. For each species alignment we removed base pair ambiguities (M, N, R, W or Y). While many authors delete singletons as they may potentially represent post-mortem damage^47^, it has been demonstrated that this ultraconservative practice can erase valuable information in the alignments and lead to biased results^48^, thus, we retained singletons in the alignments. A common issue when working with publicly available genetic data is the large variation in the length of the sequences. Therefore, we trimmed the sequences to the length of the majority of the sequences in the alignment (51%). Some aDNA sequences are typically very short (< 100 base pairs, bp), but they still usually contain valuable genetic variation for understanding temporal patterns of genetic change. Here, we retained short aDNA sequences in the alignments due to the limited availability of aDNA, as suggested in other studies^49^. Specific sites in the alignment were removed if more than 50% of the sequences had gaps or unresolved nucleotides. Finally, if a sequence did not overlap with other sequences, it was removed from the alignment.

### Genetic diversity change

To compare the global Late Pleistocene and Holocene nucleotide diversity, we categorized sequences for each species as either Late Pleistocene or Holocene based on whether the mean calibrated age of the sequence fell before or after 11,700 years BP. A separate MUSCLE alignment was performed for those two periods and genetic diversity was estimated as the nucleotide diversity (the sum of the number of differences between pairs of sequences divided by the number of pairwise comparisons)^50^ and haplotype diversity (based on haplotype frequencies)^51^ in each period using the “ape” package in R^52^. Similarly, to estimate the change in genetic diversity before and after the fastest and slowest changes in climate change velocity, we selected sequences within 5,000 years before and after the selected time period (Supplementary Table 3). Bins of 5,000 years allowed us to include enough sequences for the estimation of genetic diversity while avoiding overlap of fast and slow events (Supplementary Table 4).

Correcting for the potential effects of heterochrony (differences in sampling time of the sequences) when estimating nucleotide diversity has been recommended by some authors, who argue that not correcting for heterochrony could lead to an inflation of several summary statistics due to increased coalescence times and subsequent overestimation of polymorphisms^53, 54^. However, we did not apply this correction because 1) samples collected over short temporal extents relative to an evolutionary time span should suffer only a small inflation of the summary statistics^54^, and 2) existing methods for calculating the correction have been criticized for their circularity, potentially biasing estimates in the other direction^53^. Thus, we followed the traditional method for estimating ND to avoid extra estimation bias.

The fossil record, and by consequence aDNA, is not uniformly distributed through time, as the preservation of DNA-bearing remains depends on local conditions such as temperature and humidity. To explore whether the differences in ND were influenced by the differences in sample size between these two periods, we subsampled with replacement the period with more sequences to match the number of sequences in the period with fewer sequences. We then repeated the analyses underlying Figure 3, using the mean from the 1000 permutations of ND rather than the empirical estimate (Supplementary Fig. 1, 2 and 3).

Finally, we tested whether variation in the estimated changes in nucleotide and haplotype diversity across species can be explained by climate change velocity. We used the fastest and slowest median climate change velocity experienced by each species (Fig. 1b) as independent variable and the associated changes in genetic diversity (see above) as response variable and built simple linear models. Results are given in Fig. 4.

## Data availability

The data that support the findings of this study are available from several databases listed in the Methods of the manuscript. Data are available from the authors on reasonable request.

## Code availability

The main R functions and packages used in this study are provided in the Methods. Full R scripts are available from the authors upon request.

## Supplementary information

**Supplementary Fig. 1.**
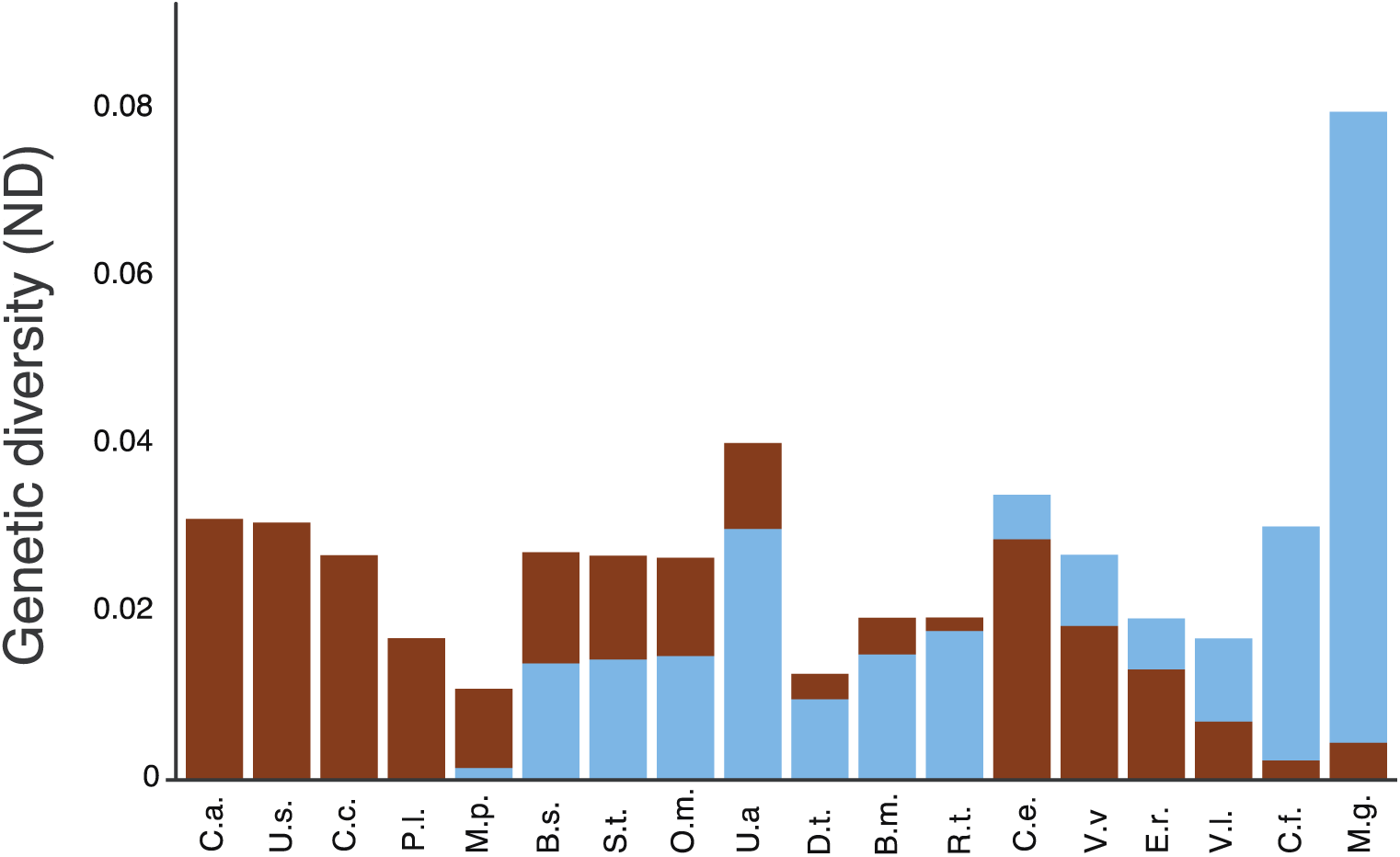
Pleistocene and Holocene genetic diversity per species. Reconstruction of Figure 3, where nucleotide diversity (ND) is the mean value of the 1000 subsampled ND estimates, so that late Pleistocene (brown) and Holocene (blue) estimates are based on the same number of sequences. Abbreviations as in Figure 1.

**Supplementary Fig. 2.**
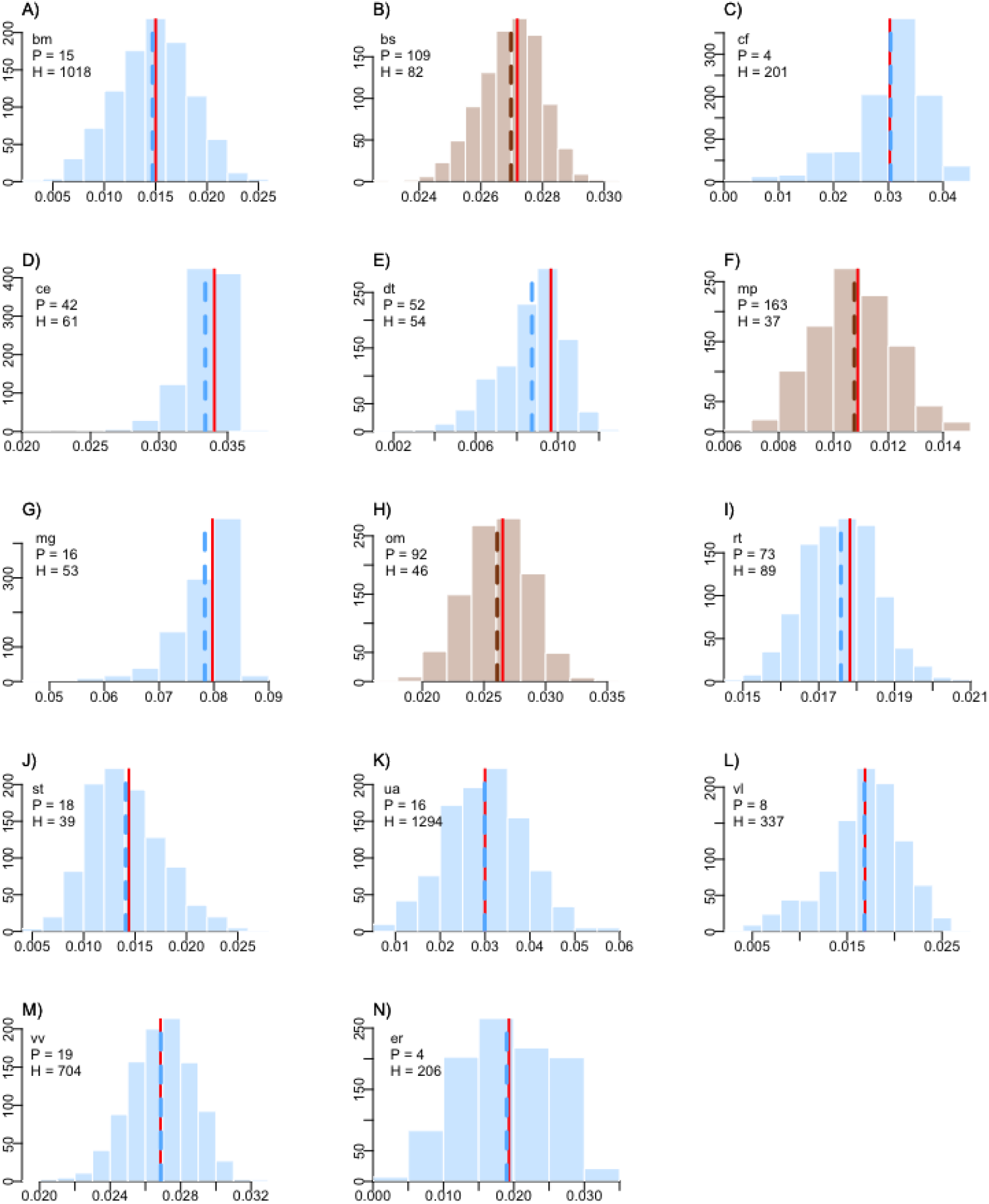
Effect of DNA sequence number in estimates of genetic diversity. Distribution of 1000 estimates of Nucleotide Diversity (ND) for the late Pleistocene and Holocene (before and after 11,700 y BP), subsampling the larger sample set to the same sample size as the period with fewer samples. Bars indicate the distribution of ND after 1000 permutations; blue distributions correspond to species where the subsamples were taken from the Holocene while brown distributions correspond to species were Pleistocene samples were subsampled. The blue or brown dashed vertical line represents the mean ND for the 1000 subsampled estimations. Red vertical line represents the ND estimation using all sequences in the database. Species abbreviations (top left in each subplot) as in Figure 1.

**Supplementary Fig. 3.**
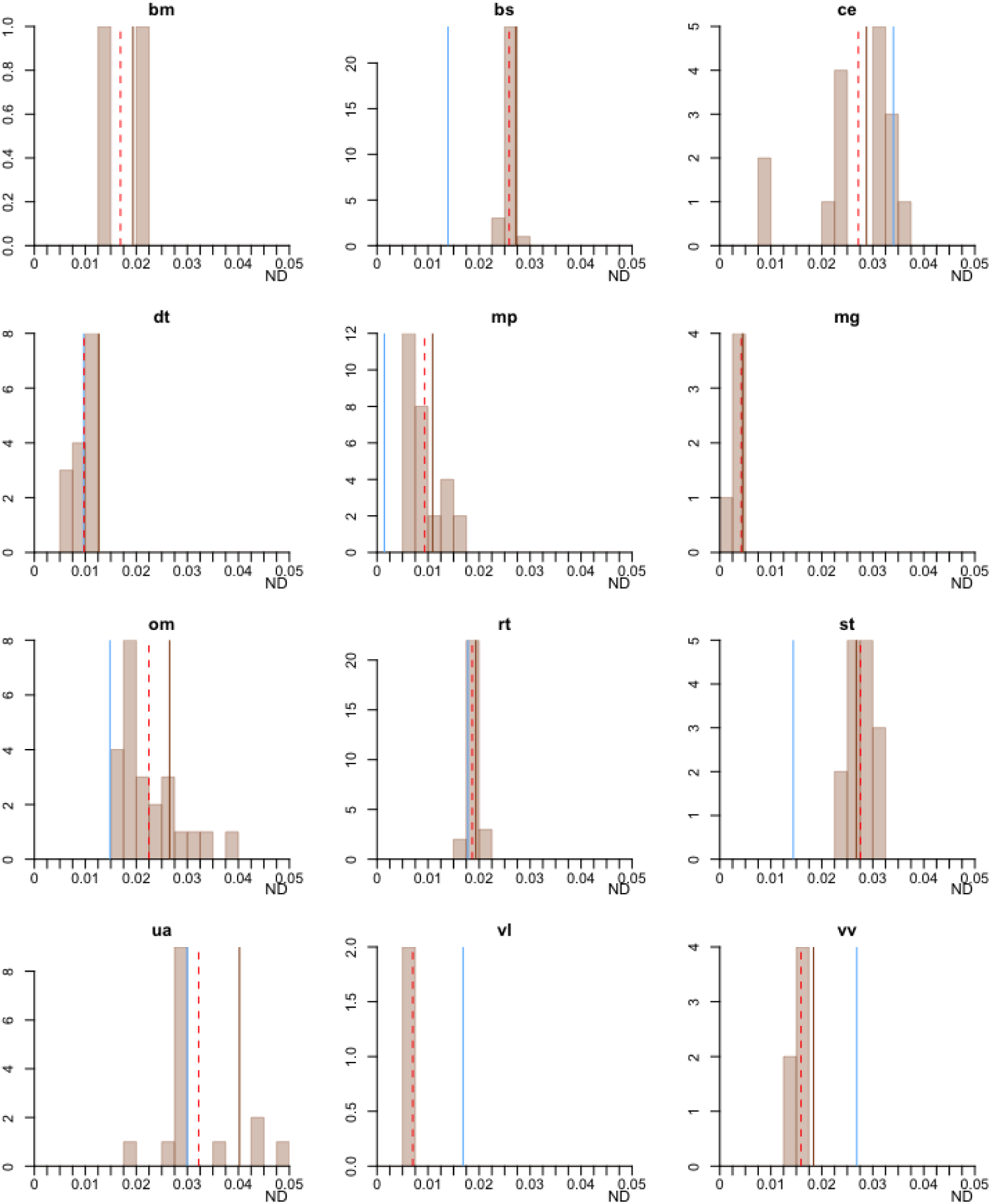
Effect of the length of DNA sequence time span in estimates of genetic diversity. Distribution of 1000 estimates of Nucleotide Diversity (ND) for the late Pleistocene and Holocene (before and after 11,700 y BP), subsampling the larger sample set to the same sample size as the period with fewer samples. Here we used a window of 11,700 years for the Pleistocene period so that the time span of the sequences sampled is the same for both periods. Bars indicate the distribution of ND after 1000 permutations; blue distributions correspond to species where the subsamples were taken from the Holocene while brown distributions correspond to species were Pleistocene samples were subsampled. The blue or brown vertical line represents the mean ND for the 1000 subsampled estimations. Red dashed vertical line represents the ND estimation using all sequences in the database. Species abbreviations (top of each subplot) as in Figure 1.

**Supplementary Fig. 4.**
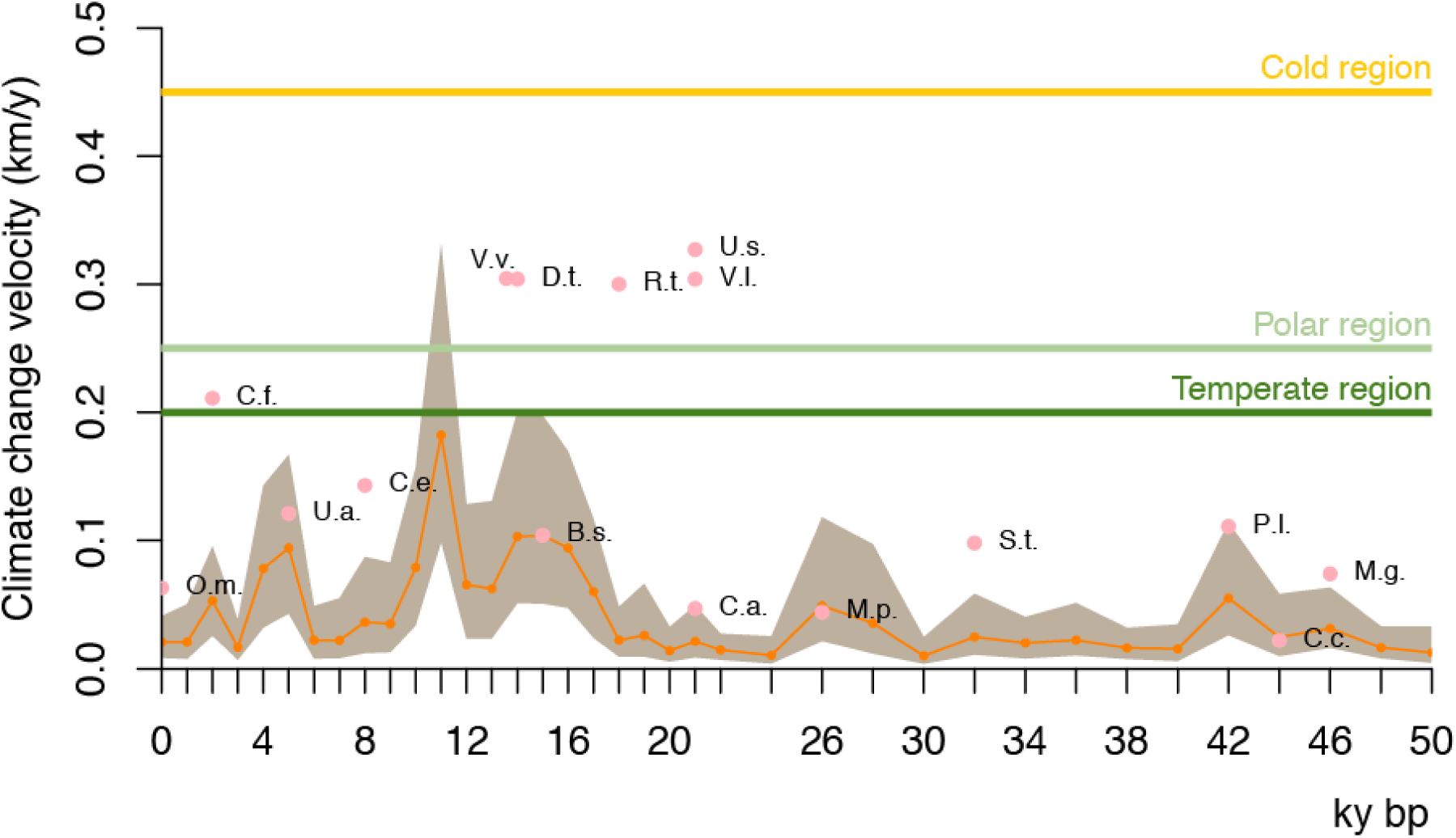
Variation in climate change velocity for the Holarctic region. Orange dots indicate the median velocity per period, while the brown area delineates the 25th and 75th percentiles. Pink points illustrate the period in which each of the 16 terrestrial species experienced the fastest climate change during the last 50,000 years. Horizontal lines indicate predicted median estimates of climate change velocity for the year 2100 for the temperate, cold and polar regions^17^.

**Supplementary Fig. 5.**
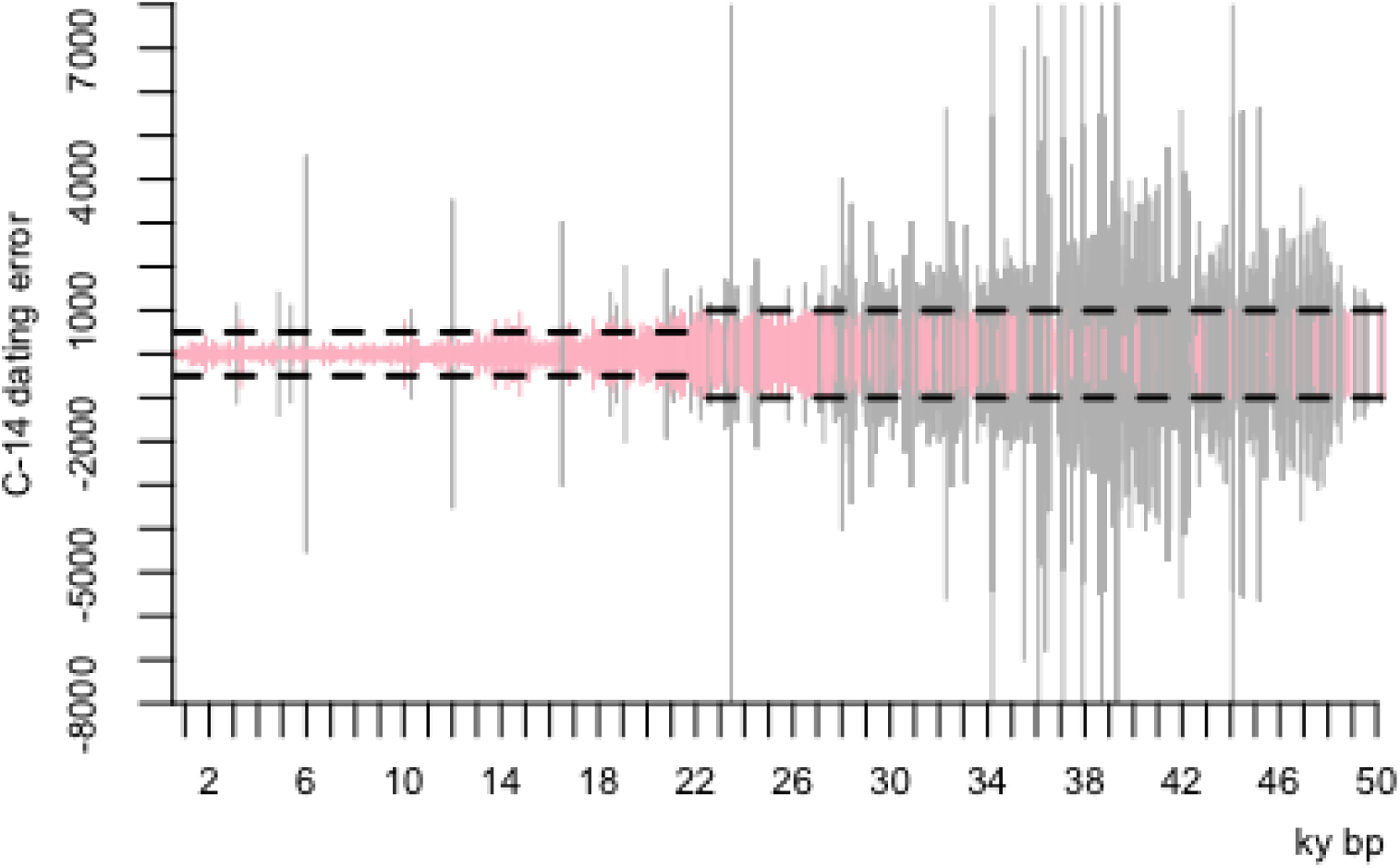
Median C^14^ dates and dating error for all fossils and aDNA sequences. The x-axis indicates median date for each sample (in kyr BP), while the y-axis indicates the dating error, shown for each sample as the height of the vertical line. Pink lines represent samples with error ≤ 1000 years. Grey lines represent samples with dating error larger than 1000 years. Horizontal dashed lines indicate the temporal window of the C^14^ dating error for samples included in each time bin for the climate change velocity analyses.

**Supplementary Table 1.** Fossil and DNA data for 18 Holarctic mammals. Total records include fossil and DNA localities. For total DNA, sequences dated older than 11,700 years are shown in the lpDNA column and sampling for Holocene sequences is shown in hDNA column. Species with names in bold are extinct.

| Species | Total records | Fossils | Total DNA | lpDNA | hDNA |
| --- | --- | --- | --- | --- | --- |
| *Balaena mysticetus* | 989 | 0 | 989 | 24 | 965 |
| *Bison sp* | 539 | 348 | 191 | 109 | 82 |
| *Castor fiber* | 288 | 88 | 200 | 4 | 196 |
| *Cervus elaphus* | 636 | 533 | 103 | 42 | 61 |
| ***Coelodonta antiquitatis*** | 535 | 480 | 55 | 55 | 0 |
| *Crocota crocuta* | 268 | 256 | 12 | 12 | 0 |
| *Dicrostonyx torquatus* | 187 | 81 | 106 | 52 | 54 |
| *Eschrichtius robustus* | 81 | 0 | 81 | 4 | 77 |
| ***Mammuthus primigenius*** | 1867 | 1664 | 203 | 165 | 38 |
| *Microtus gregalis* | 152 | 83 | 69 | 16 | 53 |
| *Ovibos moschatus* | 266 | 128 | 138 | 92 | 46 |
| *Panthera leo* | 342 | 319 | 23 | 23 | 0 |
| *Rangifer tarandus* | 853 | 691 | 162 | 73 | 89 |
| *Saiga tatarica* | 90 | 33 | 57 | 18 | 39 |
| *Ursus arctos* | 1532 | 195 | 1337 | 40 | 1297 |
| ***Ursus spelaeus*** | 386 | 386 | 0 | 0 | 0 |
| *Vulpes lagopus* | 666 | 321 | 345 | 8 | 337 |
| *Vulpes vulpes* | 1157 | 434 | 723 | 19 | 704 |
| Total | 10834 | 6040 | 4794 | 756 | 4038 |

**Supplementary Table 2.**
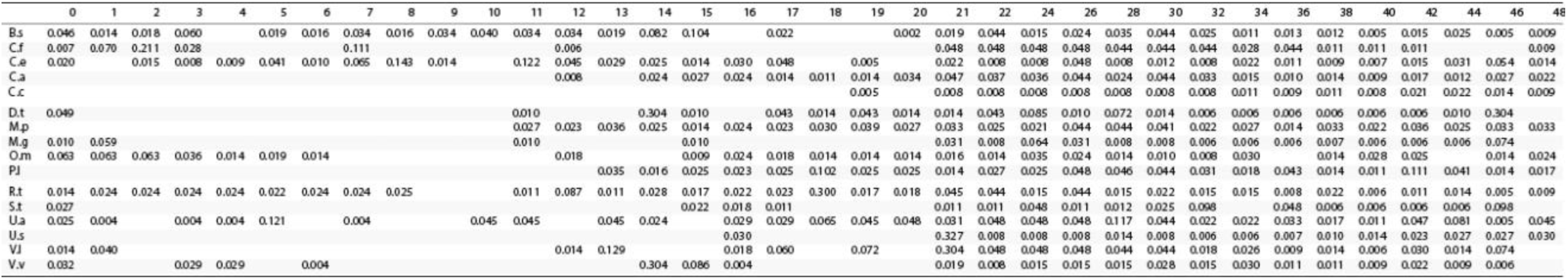
Median climate change velocity per time bin for 16 terrestrial mammals. Columns indicate the upper (younger) bound of the time bin in thousands of years before present. For example, column 48 shows the median climate change velocity from 50,000 to 48,000 y BP. Species abbreviations as in Figure 1.

**Supplementary Table 3.** Time periods (in years before present) with the fastest and slowest climate change velocity per species. Fast and slow columns represent the time periods with enough sequences before and after the change in climate to test for changes in genetic diversity.

| Species | Fastest | Slowest | Fast | Slow |
| --- | --- | --- | --- | --- |
| *Bison sp* | 15000 | 20000 | 15000 | 40000 |
| *Castor fiber* | 2000 | 12000 | 2000 | 12000 |
| *Cervus elaphus* | 8000 | 19000 | 11000 | 3000 |
| *Coelodonta antiquitatis* | 21000 | 12000 | 21000 | 40000 |
| *Crocota crocuta* | 44000 | 19000 | 44000 |  |
| *Dicrostonyx torquatus* | 14000 | 32000 | 14000 | 26000 |
| *Mammuthus primigenius* | 26000 | 15000 | 26000 | 15000 |
| *Microtus gregalis* | 46000 | 32000 |  | 15000 |
| *Ovibos moschatus* | 0 | 32000 | 1000 | 18000 |
| *Panthera leo* | 42000 | 40000 | 18000 |  |
| *Rangifer tarandus* | 18000 | 46000 | 18000 | 36000 |
| *Saiga tatarica* | 32000 | 38000 |  | 17000 |
| *Ursus arctos* | 5000 | 1000 | 18000 | 1000 |
| *Ursus spelaeus* | 21000 | 32000 |  |  |
| *Vulpes lagopus* | 21000 | 40000 |  |  |
| *Vulpes vulpes* | 14000 | 6000 | 15000 | 6000 |

**Supplementary Table 4.** DNA sequences used for associations between climate change velocity and genetic diversity change. Available per species for 3,000, 5,000 and 7,000 years before and after the time period with the fastest and slowest climate change velocity (see Supplementary Table 3).

| Species | Fast |  |  |  |  |  | Slow |  |  |  |  |  |
| --- | --- | --- | --- | --- | --- | --- | --- | --- | --- | --- | --- | --- |
|  | 3000 |  | 5000 |  | 7000 |  | 3000 |  | 5000 |  | 7000 |  |
|  | After | Before | After | Before | After | Before | After | Before | After | Before | After | Before |
| Bison sp | 20 | 11 | 25 | 11 | 28 | 14 | 1 | 3 | 11 | 12 | 24 | 18 |
| Castor fiber | 168 | 10 | 168 | 12 | 168 | 25 | 3 | 4 | 16 | 4 | 18 | 4 |
| Cervus elaphus | 4 | 2 | 7 | 6 | 8 | 18 | 4 | 1 | 12 | 2 | 19 | 2 |
| Coelodonta antiquitatis | 5 | 5 | 10 | 13 | 15 | 14 |  | 2 |  | 7 |  | 13 |
| Crocota crocuta | 2 | 4 | 5 | 7 | 5 | 7 |  |  |  |  |  |  |
| Dicrostonyx torquatus | 21 | 20 | 21 | 22 | 21 | 25 | 3 |  | 3 |  | 19 |  |
| Mammuthus primigenius | 12 | 12 | 18 | 23 | 23 | 33 | 10 | 11 | 12 | 21 | 19 | 28 |
| Microtus gregalis |  |  |  |  |  |  |  |  |  |  |  |  |
| Ovibos moschatus |  | 29 |  | 44 |  | 46 | 7 |  | 16 |  | 32 | 1 |
| Panthera leo |  |  |  |  | 1 | 1 |  |  | 1 |  | 1 |  |
| Rangifer tarandus | 11 | 8 | 13 | 16 | 15 | 21 |  |  | 1 |  | 2 |  |
| Saiga tatarica | 3 |  | 3 |  | 5 |  |  |  |  |  | 1 |  |
| Ursus arctos | 11 | 11 | 1275 | 15 | 1275 | 22 | 1246 | 25 | 1246 | 34 | 1246 | 40 |
| Ursus spelaeus |  |  |  |  |  |  |  |  |  |  |  |  |
| Vulpes lagopus |  |  | 2 |  | 2 |  |  |  |  |  |  |  |
| Vulpes vulpes | 2 | 10 | 2 | 10 | 2 | 10 | 3 | 8 | 4 | 8 | 696 | 10 |

## References

1. Koch, P. L. & Barnosky, A. D. Late Quaternary Extinctions: State of the Debate. Annu. Rev. Ecol. Evol. Syst. 37, 215–250 (2006).

2. Blois, J. L., McGuire, J. L. & Hadly, E. A. Small mammal diversity loss in response to late-Pleistocene climatic change. Nature 465, 771–774 (2010).

3. Cooper, A. et al. Abrupt warming events drove Late Pleistocene Holarctic megafaunal turnover. Science. 349, 602–606 (2015).

4. Rabanus-Wallace, M. T. et al. Megafaunal isotopes reveal role of increased moisture on rangeland during late Pleistocene extinctions. Nat. Ecol. Evol. 1, 1–5 (2017).

5. Lorenzen, E. D. et al. Species-specific responses of Late Quaternary megafauna to climate and humans. Nature 479, 359–364 (2011).

6. Palkopoulou, E. et al. Complete genomes reveal signatures of demographic and genetic declines in the woolly mammoth. Curr. Biol. 25, 1395–1400 (2015).

7. Leonardi, M. et al. Evolutionary patterns and processes: lessons from ancient DNA. Syst. Biol. 66, e1–e29 (2017).

8. Barnosky, A. D. et al. Has the Earth/’s sixth mass extinction already arrived? Nature 471, 51–57 (2011).

9. Spielman, D., Brook, B. W. & Frankham, R. Most species are not driven to extinction before genetic factors impact them. Proc. Natl. Acad. Sci. U. S. A. 101, 15261–15264 (2004).

10. Mimura, M., et al. Understanding and monitoring the consequences of human impacts on intraspecific variation. Evol. Appl. 10, 121–139 (2017).

11. Oliver, T. H. et al. Biodiversity and resilience of ecosystem functions. Trends Ecol. Evol. 30, 673–684 (2015).

12. Barnosky, A. D. et al. Merging paleobiology with conservation biology to guide the future of terrestrial ecosystems. Science 355, (2017).

13. Zheng, J., Payne, J. L., Wagner, A. Cryptic genetic variation accelerates evolution by opening access to diverse adaptive peaks. Science 365, 347–353 (2019).

14. Hoffmann, A. A. & Sgro, C. M. Climate change and evolutionary adaptation. Nature 470, 479–485 (2011)

15. Campos, P. F. et al. Ancient DNA analyses exclude humans as the driving force behind late Pleistocene musk ox (Ovibos moschatus) population dynamics. Proc. Natl. Acad. Sci. 107, 5675–5680 (2010).

16. Singarayer, J. S., Valdes, P. J., Friedlingstein, P. & Nelson, S. Late Holocene methane rise caused by orbitally controlled increase in tropical sources. Nature (2011)

17. Garcia, R. A., Cabeza, M., Rahbek, C. & Araújo, M. B. Multiple dimensions of climate change and their implications for biodiversity. Science. 344, 1247579 (2014).

18. Graham, R. W. et al. Spatial Response of Mammals to Late Quaternary Environmental Fluctuations. Science. 272, 1601–1606 (1996).

19. Willerslev, E. et al. Fifty thousand years of Arctic vegetation and megafaunal diet. Nature 506, 47–51 (2014).

20. Stewart, L., Alsos, I. G., Bay, C. & Breen, A. L. The regional species richness and genetic diversity of Arctic vegetation reflect both past glaciations and current climate. Glob. Ecol. Biogeogr. 25, 430–442 (2016).

21. Vellend, M. & Geber, M. A. Connections between species diversity and genetic diversity. Ecol. Lett. 8, 767–781 (2005).

22. Orlando, L. & Cooper, A. Using ancient DNA to understand evolutionary and ecological processes. Annu. Rev. Ecol. 44, 573–598 (2014).

23. Romiguier, J. et al. Comparative population genomics in animals uncovers the determinants of genetic diversity. Nature 515, 261–263 (2014).

24. Nogués-Bravo, D., Rodríguez, J., Hortal, J., Batra, P. & Araújo, M. B. Climate Change, Humans, and the Extinction of the Woolly Mammoth. PLoS Biol. 6, e79– e79 (2008).

25. Sandel, B. et al. The influence of Late Quaternary climate-change velocity on species endemism. Science. 334, 660–664 (2011)

26. Stewart, J. R., Lister, A. M., Barnes, I. & Dalen, L. Refugia revisited: individualistic responses of species in space and time. Proc. R. Soc. B Biol. Sci. 277, 661–671 (2010).

27. Loarie, S. R. et al. The velocity of climate change. Nature 462, 1052–1055 (2009).

28. Steffensen, J. P. et al. High-resolution Greenland ice core data show abrupt climate change happens in few years. Science. 321, 680–684 (2008).

29. Schipper, J. et al. The Status of the World’s Land and Marine Mammals: Diversity, Threat, and Knowledge. Science. 322, 225–230 (2008).

30. Barnosky, A. D. et al. Approaching a state shift in Earth’s biosphere. Nature 486, 52–58 (2012).

31. van Andel, T. H. The Climate and Landscape of the Middle Part of the Weichselian Glaciation in Europe: The Stage 3 Project. Quat. Res. 57, 2–8 (2002).

32. Ramsey, C. B., Bayesian Analysis of Radiocarbon Dates. Radiocarbon. 51, 337– 360 (2009).

33. Ramsey, C. B., Lee, S., Recent and Planned Developments of the Program OxCal. Radiocarbon. 55, 720–730 (2013).

34. Reimer, P., IntCal13 and Marine13 Radiocarbon Age Calibration Curves 0–50,000 Years cal BP. Radiocarbon. 55, 1869–1887 (2013).

35. Wick, M., Vatant, B., The geonames geographical database. Available from World Wide Web http://geonames.org (2012).

36. Miraldo, A., et al. An Anthropocene map of genetic diversity. Science. (2016)

37. Gordon C., et al. The simulation of SST, sea ice extents and ocean heat transports in a version of the Hadley Centre coupled model without flux adjustments. Clim. Dyn. 16, 147–168 (2000).

38. Varela, S., Lima-Ribeiro, M. S., Diniz-Filho, J. a F., Storch, D. Differential effects of temperature change and human impact on European Late Quaternary mammalian extinctions. Glob. Chang. Biol. 1–7 (2014).

39. Singarayer, J. S., Valdes, P. J. High-latitude climate sensitivity to ice-sheet forcing over the last 120kyr. Quat. Sci. Rev. (2010).

40. Diaz, J. M. S., Franklin, J., Ninyerola, M. Bioclimatic velocity: the pace of species exposure to climate change. Divers. Distrib. (2014).

41. Brito-Morales I., et al. Climate Velocity Can Inform Conservation in a Warming World. Trends Ecol. Evol. (2018).

42. Dobrowski, S. Z., Parks, S. A. Climate change velocity underestimates climate change exposure in mountainous regions. Nat. Commun. (2016).

43. R Core Team, R: A Language and Environment for Statistical Computing. R Found. Stat. Comput. Vienna Austria. 0 (2014), p. {ISBN} 3-900051-07-0,, doi:10.1017/CBO9781107415324.004.

44. Hijmans, R. J., van Etten, J. raster: Geographic analysis and modeling with raster data. R Packag. version 2.5-2 (2015), p. http://CRAN.R-project.org/package=raster.

45. Edgar, R. C., MUSCLE: multiple sequence alignment with high accuracy and high throughput. Nucleic Acids Res. 32, 1792–1797 (2004).

46. Kearse, M., et al. Geneious Basic: An integrated and extendable desktop software platform for the organization and analysis of sequence data. Bioinformatics. 28, 1647–1649 (2012).

47. Axelsson, E., Willerslev, E., Gilbert, M. T. P., Nielsen, R. The Effect of Ancient DNA Damage on Inferences of Demographic Histories. Mol. Biol. Evol. 25, 2181– 2187 (2008).

48. Rambaut, A., Ho, S. Y. W. Drummond, A. J., Shapiro, B., Accommodating the Effect of Ancient DNA Damage on Inferences of Demographic Histories. Mol. Biol. Evol. 26, 245–248 (2009).

49. Palkopoulou, E., et al. Holarctic genetic structure and range dynamics in the woolly mammoth. Proc. R. Soc. B Biol. Sci. 280, 20131910 (2013).

50. Nei, M., Li, W. H. Mathematical model for studying genetic variation in terms of restriction endonucleases. Proc. Natl. Acad. Sci. (1979). E. Paradis, J. Claude, K. Strimmer, APE: Analyses of Phylogenetics and Evolution in R language. Bioinformatics. 20, 289–290 (2004).

51. Nei, M., Tajima, F. DNA polymorphism detectable by restriction endonucleases. Genetics (1981). doi:10.1016/j.chemphyslip.2007.10.006

52. Navascués, M., Depaulis, F., Emerson, B. C., Combining contemporary and ancient DNA in population genetic and phylogeographical studies. Mol. Ecol. Resour. 10, 760–772 (2010).

53. Depaulis, F., Orlando, L., Hänni, C. Using classical population genetics tools with heterochroneous data: time matters! PLoS One. 4, e5541 (2009).

54. Welch, A. J., et al. in Molecular Biology and Evolution (2012), vol. 29, pp. 3729–3740.

